# Parallel progress in perceived age and life expectancy

**DOI:** 10.1101/195990

**Authors:** Ulrich K. Steiner, Lisbeth Aagaard Larsen, Kaare Christensen

## Abstract

Human life expectancy continues to rise in most populations. This rise not only leads to longer lives but is also accompanied by improved health at a given age, i.e. we see a reduction of biological age for a given chronological age in recent cohorts. Despite or even because of the diversity of biomarkers of aging, an accurate quantification of a general shift in biological age across time has been challenging. By comparing age perception of images taken in 2001 over a decade, we show that age perception changes substantially across time and parallels the progress in life expectancy. In 2012, people aged 70+ needed to look 2.3 years younger to be rated the same age as in 2002. Our results further suggest that age perception reflects the past life events better than predicts future length of life, i.e. it is written in your face how much you have aged so far, but does not predict well how fast you will age in the future. We draw this conclusion since age perception among elderly paralleled changes in life expectancy at birth but not changes in remaining life expectancies. We illustrate advantages of perceived age as a biomarker of aging and suggest that changes in age perception should be explored for younger age classes to inform on aging processes, including whether aging is delayed or slowed with increasing life expectancy.

## Introduction

Human life expectancy has risen in long-lived populations by about two years per decade (Oeppen & Vaupel 2002; Vaupel 2010). This continuous and exceptional rise in lifespan started around one and a half centuries ago and not only lead to longer lives, but has also been accompanied by increased healthspan (Christensen, Doblhammer, et al. 2009; Crimmins & Beltran-Sanchez 2011). That is, with increasing life expectancy we observe improved health and a reduction in biological age measured by improved scores on biomarkers of health and aging at a given chronological age. Middle-aged people nowadays show improved health compared to former times (Gerstorf et al. 2011), and improvements are not limited to younger people or the most recent cohorts. Even the oldest old (>90 years) score better on biomarkers of aging when born in more recent cohorts compared to earlier cohorts (Christensen et al. 2013). Such findings of shifting biological age indicate that people in recent times i) age less fast, ii) delay aging, or iii) begin life at a lower level of biological age.

Despite the qualitative evidence for prolonged healthy lives and improved physiological and cognitive function of recent birth cohorts, quantifying the relationship between rising life expectancy and improved biological age has been challenging (Crimmins & Vasunilashorn 2011; Crimmins et al. 2005). Biomarkers of aging provide the main tool to investigate biological age but are often used in a qualitative rather than quantitative way. Many biomarkers identify threshold levels that distinguish different groups of biological age rather than providing continuous measures of biological age (Mitnitski et al. 2015). This categorization in groups relates to another frequent challenge in evaluating biomarkers: how to convert the unit a biomarker is measured in —microunits per millilitre for insulin, kilograms for grip strength, or some scoring of cognitive tests — to changes in hazard risk, related mortality, and consequently life expectancy. In consequence, relating the biological age to the biological process of aging as aimed at with hundreds of different biomarkers remains difficult. To define the underlying aging process in relationship to the biological and chronological age, one also needs to account for partial correlation between average chronological aging processes, the interplay among the different biomarkers, and the biomarker environment interaction. The challenge of how to best evaluate biological age is also illustrated by the many different (healthy) aging indices that have been developed, and the controversy surrounding how to weigh the contributing biomarkers (Sanders et al. 2014; Tyrovolas et al. 2014; Forti et al. 2012; Levine 2013; Mitnitski et al. 2015). To this end it is not clear how we can derive a simple and meaningful integrated measure of biological age.

One of the many biomarkers of aging is perceived age, i.e. how old one looks, which is associated with survival, and physical and cognitive functioning (Christensen, Thinggaard, et al. 2009; Dykiert et al. 2012). For people aged 70 and older, even after correcting for sex and chronological age as well as other partly correlated biomarkers — e.g. cognitive scores, strength scores, grip strength, and the mini-mental state examination — annual mortality risk increased by 3% with each year deviation in perceived age from chronological age (Christensen, Thinggaard, et al. 2009). Younger people aged 30-70, had relatively bad health if their perceived age was judged substantially older than their chronological age, and perceived age correlated with other biomarkers of aging in a follow up study comparing the same individuals at age 26 and 38 (Hwang et al. 2011; Belsky et al. 2015). Perceived age might therefore be seen as an integrative and relatively general biomarker of aging with a major advantage compared to many other biomarkers: both perceived age and chronological age are measured in the same units, age in years, and can therefore be directly compared to increased life expectancy.

Here we used the change in perceived age of the same facial images assessed over a decade to evaluate how this measure of biological age changed over that decade and how this change relates to shifts in cohort life expectancy. For our evaluation we used facial images taken in 2001 (February-April) and collected two rounds of perceived age on these images; the first round was collected in November 2002 and the second round was collected in December 2012. Obviously, the images have not changed over the ten-year period between the round of assessments. The naïve expectation would be that the perceived age should not change, which would indicate that the biological age — measured as perceived age — did not change over a decade. However, as we argue here, if perceived age tracks the change in biological age across time we expect that the same images evaluated at the end of 2002 would be judged to be younger and closer to their chronological age compared to the evaluation ten years later. Our aim was to test the hypothesis that the difference in perceived age between the two assessments quantifies the change in biological age and parallels the process of changed cohort life expectancy over this decade.

## Results

Mean chronological age of the 238 Danish women, and 144 Danish men aged 70+ of whose facial images were taken in 2001 was 75.9 years. Both sexes showed similar chronological age distributions (males mean 75.5; females mean 76.2; Null model: no difference between sexes AIC=4512.4, Model with sex differences AIC=4510.7) (Table. 1; Fig. S1). Perceived mean age in 2002 and 2012 was lower for males than for females (Table. 1; Fig. S1 & Table S1). The difference between the sexes in perceived age was about equal between 2002 and 2012; at least it did not differ substantially among the assessment years (similar support [δAIC<2] for an additive model compared to an interactive model between sex and assessment year; Table S1). The distribution of chronological age of the sample images did not differ among the sexes, but perceived age differed between males and females and between the assessment years.

**Table 1:** Mean and standard deviation of chronological and perceived age, deviance between chronological and perceived age, and change in life expectancies in Denmark. All measurements are in years.

| Parameters | Mean±SD (Minimum, Maximum) |  |  |
| --- | --- | --- | --- |
|  | Both sexes | Males | Females |
| Chronological Age | 75.9±4.6 (70,90) | 75.5±4.8 (70,90) | 76.2±4.5 (70,89) |
| Perceived Age 2002 | 76.5±3.9 (65,90) | 76.3±3.4 (65,86) | 76.6±4.2 (65,86) |
| Deviance between perceived age 2002 and chronological age | 0.6 ±5.0 (-16, 11) | 0.8 ±5.3 (-16, 11) | 0.4±4.9 (-14, 11) |
| Perceived Age 2012 | 78.8±4.2 (68,90) | 78.1±3.6 (68,88) | 79.2±4.5 (68,91) |
| Deviance between perceived age 2012 and chronological age | 2.9 ±3.9 (-9, 14) | 2.6 ± 4.3 (-8, 12) | 3.0 ±4.4 (-9, 14) |
| period life expectancy 2003* | 77.7 | 75.3 | 80.1 |
| period life expectancy 2013* | 80.3 | 78.3 | 82.3 |
| Change in period life expectancy 2003-2013 | 2.6 | 3.0 | 2.25 |
| 1913 cohort life expectancy | 64.9 | 61.8 | 68.1 |
| 1923 cohort life expectancy | 67.2 | 63.9 | 70.6 |
| Change in cohort life expectancy (1913 to 1923 cohort) | 2.3 | 2.1 | 2.5 |

\* closest period estimates (Jan. 2003 & Jan. 2013) to the perceived age estimates (November 2002 and December 2012) (Wilmoth & Shkolnikov 2011).

Mean perceived age in November 2002 was rated to be about half a year higher than the chronological age of the images taken in spring of 2001, while mean perceived age in December 2012 was rated to be almost three years older than the chronological age and 2.3 years older compared to the 2002 assessment (Table 1). The deviance between chronological and perceived age did not differ substantially among the sexes, either in 2002 or in 2012 (Table 1&2, Fig. 1). To this end, perception of age changed substantially over a decade in males and females to similar extents (Table 1&2).

**Fig. 1.**
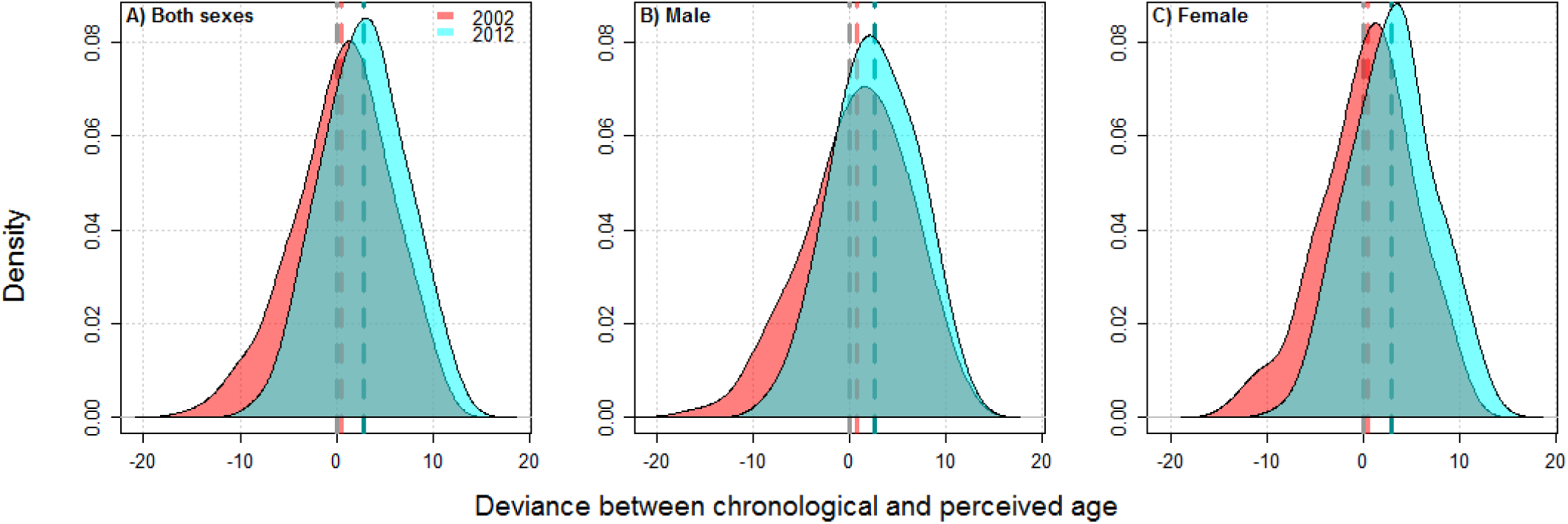
Smoothed distribution of deviance between chronological and perceived age in 2002 (red) and 2012 (blue) for A) both sexes combined, B) males, and C) females. Positive deviances show higher perceived age compared to the chronological age while negative values show the opposite. The vertical grey dashed line marks zero deviance, i.e. the exact mean chronological age, the dashed red line marks the mean deviance between the chronological age and the perceived age in 2002, and the blue hatched line marks the mean deviance between the chronological age and the perceived age in 2012.

**Table 2:** Deviance between chronological and perceived age as a function of sex, rating year (Year), and chronological age. Slopes and quadratic terms are only reported for the continuous factor chronological age. The main mean differences are reported in the main text and illustrated in Fig. 1, Fig. S1, Fig.2. Lower AIC values indicate better statistical support for the model. Bold AIC values highlight the best supported model with one, two, or three (or more) factors, respectively.

| Factors | Slope±SE | Quadratic±SE term | df | AIC |
| --- | --- | --- | --- | --- |
| Null model (intercept only) | - | - | 2 | 4579.11 |
| Sex | - | - | 3 | 4581.10 |
| Year | - | - | 3 | **4535.67** |
| Sex+Year (additive) | - | - | 4 | 4537.66 |
| Sex*Year (interactive) | - | - | 5 | 4538.55 |
| Chronological Age | -0.62±0.03 | - | 3 | 4244.09 |
| Sex+Chronological Age | -0.62±0.03 | - | 4 | 4243.67 |
| Sex*Chronological Age | -0.56±0.04 (females)<br>-0.74±0.06 (males) | - | 5 | 4237.26 |
| Year+Chronological Age | -0.62±0.03 | - | 4 | 4174.25 |
| Year*Chronological Age | -0.73±0.04 (2002)<br>-0.51±0.06 (2012) | - | 5 | **4161.94** |
| Sex+Year+Chronological Age | -0.63±0.03 | - | 5 | 4173.59 |
| Sex+Year*Chronological Age | -0.74±0.04 (2002)<br>-0.52±0.06 (2012) | - | 6 | 4161.23 |
| Year+Sex*Chronological Age | -0.56±0.04 (females)<br>-0.73±0.06 (males) | - | 6 | 4166.34 |
| Sex*Year*Chronological Age | -0.65±0.05 (females 2002)<br>-0.86±0.08 (males 2002)<br>-0.46±0.07 (females 2012)<br>-0.60±0.12 (males 2012) | - | 9 | **4156.37** |
| Chronological Age+(Chron. Age)<sup>2</sup> | -0.72±0.88 | -0.0006±0.0057 | 6 | 4246.08 |
| Year*Chronological Age+(Chron. Age)<sup>2</sup> | -0.83±0.84 (2002)<br>-0.61±0.57 (2012) | -0.0006±0.0057 | 6 | 4163.93 |
| Sex*Year*Chronological Age+(Chron. Age)<sup>2</sup> | -0.95±0.84 (female 2002)<br>-1.06±0.08 (female 2002)<br>-0.76±0.07 (female 2012)<br>-0.89±0.12 (female 2012) | -0.002±0.005 | 10 | 4158.25 |

The deviation between the chronological age and the perceived age depended on the actual chronological age in males and females (Fig. 2, Table 2). Younger aged persons were more often rated to have higher perceived age than their chronological age (positive deviances), whereas older persons were more often rated to look younger than their chronological age (negative deviances). This chronological-age-specific shift was found for both males and females. The dependency on chronological age was more pronounced in 2002 than in 2012 (see Year*Chronological age estimates Table 2), and chronological age had a slightly stronger influence on how deviance changed between 2002 and 2012 in males compared to females. The relationship between the response variable (i.e. the deviance between chronological and perceived age) and chronological age was linear. Models with curvilinear terms (quadratic terms) were not better supported than such with only linear terms (Table 2). Such chronological age-dependent age perception has been frequently described before for similar data and explained as regression towards the mean (Christensen et al. 2004).

**Fig. 2:**
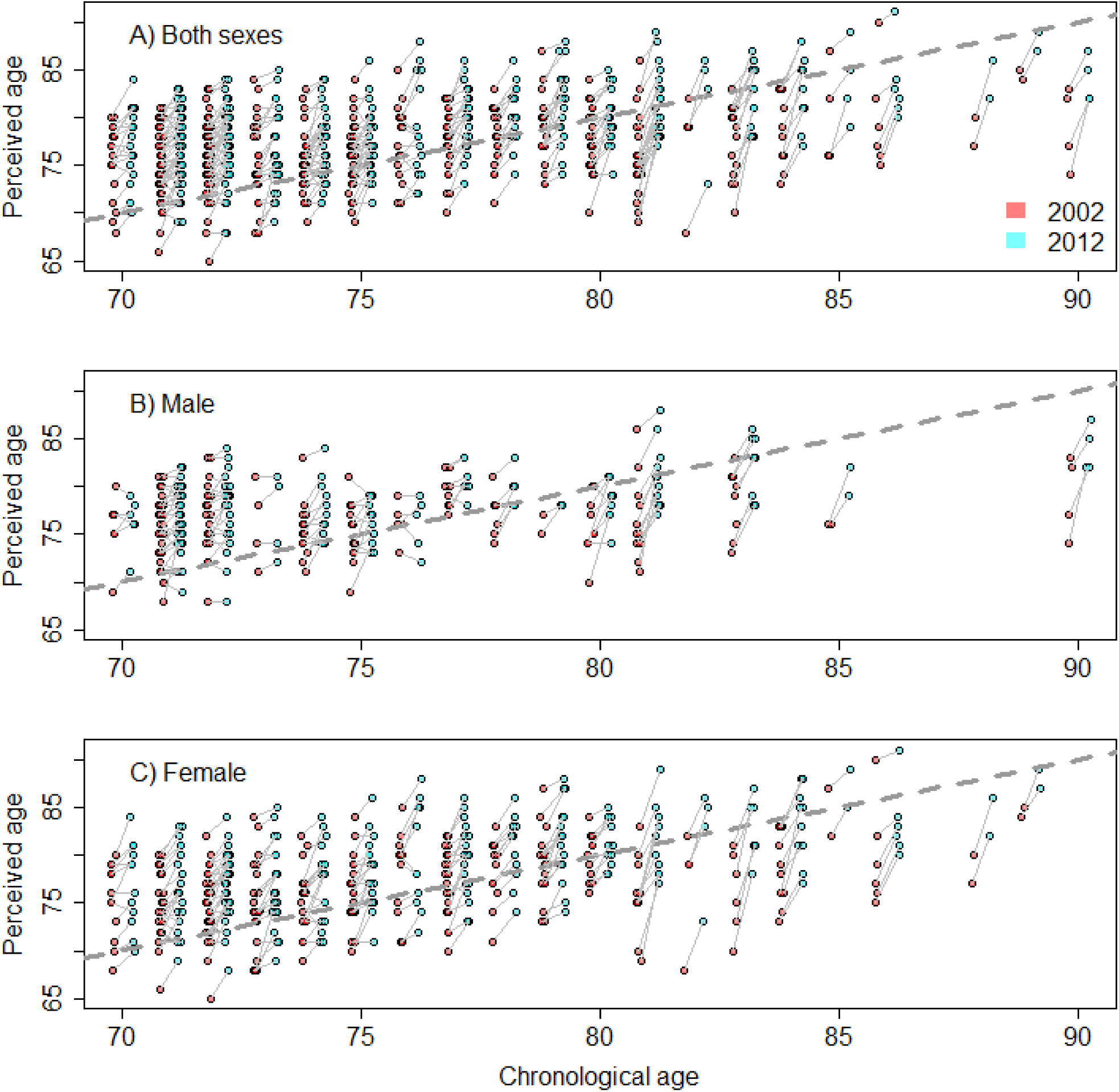
Chronological age plotted against perceived age for perceived age assessed in 2002 (red circles) and 2012 (blue circles) for both sexes combined (A), males (B) and females (C). The dashed grey lines mark the theoretical one to one chronological to perceived age relationship. The thin grey lines connect the perceived age estimates on each image rated in 2002 and 2012. For better visualization of the data a jitter and small offset between the two ratings is added to the chronological age.

Since the interpretation and visualization of three way interactions (sex*year*chronological age, Table 2) is challenging, we also illustrate the difference in the deviance between chronological and perceived age in 2002 and 2012 along the chronological age axis (Fig. S2). Such difference in the deviance corresponds to the length of the thin grey lines in Fig. 2; these lines increase in length with increasing chronological age. These differences in deviance not only depended on the chronological age, they also differed among the sexes (females mean 2.6 ±0.2; males 1.9 ±0.2), but it is less clear whether there are important interactive effects between sex and chronological age, or curvilinear (quadratic) chronological age effects (Table S2).

## Discussion

We show, as hypothesized that the perception of age changes across time, which implies that age perception adjusts to the shift in biological age, or at least the biological age to chronological age conversion changes over time. For instance, a person aged 70 in 2012 needed to look substantially younger (2.3 years) to be perceived the same age as a 70-year-old in 2002. Therefore, biological age — derived from perceived age — changed by 2.3 years over the ten-year period. This change is close to the change in life expectancy at birth in Denmark over that same period which has risen by 2.6 years (Table 1) (Wilmoth & Shkolnikov 2011). However, such similarities might not be meaningful since life expectancy at birth is a period estimate across all age classes and not only limited to older cohorts. More interesting is that cohort based life expectancy changed by similar amounts (Table 1) (Wilmoth & Shkolnikov 2011). Unfortunately cohort life expectancies for later cohorts that build the bulk of our study sample are not yet available since many members of more recent cohorts are still alive (Wilmoth & Shkolnikov 2011). Nonetheless, it is not expected that shifts in cohort life expectancy for 70+ years old over that decade would be significantly different.

Even though the shift in perceived age over the investigated decade agrees well with shifts in life expectancy, a more relevant measure for comparison might be the change in remaining cohort life expectancy, e.g. how many additional years can a 70-year-old of the 1942 cohort (being 70 in 2013) expect to live compared to a 70-year-old from the 1932 cohort. Remaining cohort life expectancy has risen between 2003 and 2013 for 70-year-old males and females by ∼0.84 years, for 80-year-old females by ∼0.72 females, and for males by ∼0.84, and for 90-year-old females by ∼0.42 females and males by ∼0.3 (Frank Hansen & Stephensen 2013). Hence, perceived age has changed much more than remaining life expectancy and is more closely associated with shifts in cohort life expectancy at birth.

Our results suggest that age perception reflects past life events better than it predicts future length of life. This implies that the past aging history of an individual, such as physiological aging processes of accumulation of oxidative damage, is better reflected in the facial features than the future potential of life, hazard risks and the resulting remaining life expectancy. Under this point of view, we might want to consider how survivorship, the probability of living to a certain age, has changed over the ten year time period. Survivorship to age 70 in 2003 (i.e. proportion of the 1932 cohort living to at least age 70) was 0.68 in females, 0.57 in males, and 0.63 when combining both sexes (Wilmoth & Shkolnikov 2011). Similar survivorships of individuals born ten years later, i.e. comparable survivorships of the 1942 cohort, were associated with 76.1 year old females, 75.6 year old males, and 75.8 year olds when both sexes are combined (polynomial projections after age 71 based on 1942 cohort death rates extracted from the Human Mortality database; Wilmoth & Shkolnikov 2011). We observe larger shifts (5.6-6.1 years) among similar survivorship levels over the ten-year period compared to the perceived age shifts. Hence, age perception does not directly relate to shifts in accumulated survival, but rather to changes in life expectancy at birth. Substantially more people reached age 70 in the 1942 cohort compared to the 1932 cohort. Our sample only included individuals aged 70+. Hence the people whose facial images were used in our study had already passed substantial parts of their life and progressed a good part along their individual aging trajectories. These past trajectories seem to leave their marks in the faces and dominate the age perception. Such findings support our argument that perceived age accurately captures the biological age of an individual as described by how far the body has declined in function over its past life.

Taking our results together, we find ourselves in a somewhat conflicting situation in that our average age rating is quite accurate and adjusts over time, but at the same time reveals an chronological age related bias. This age bias, overestimating the age of younger people and underestimating older people, could be explained by regression towards the mean, or by selective processes as related to frailty arguments, where the most frail and oldest looking individuals die first and only the most robust and young looking individuals make it to old age (Vaupel & Yashin 1985). Regression towards the mean is an effect where raters consciously or sub-consciously realize the age range of the sample images and subsequently tend to rate closer to the sample mean, i.e. they overestimate younger individuals and underestimate older individuals. If regression to the mean occurs, we expect that the variance in age perception increases with increasing numbers of images rated. At the onset of rating images the raters have not yet developed a sense of what the age range of people they rate on is, such sense will develop with increasing number of images rated upon and raters should regress more and more towards the mean. However, such pattern of convergence towards the mean with increasing number of images rated on has not been detected (Fig. S4). As for other explanations, frailty models suggest that in an aging cohort the population is composed more and more of robust individuals because the most frail individuals are more likely to die earlier in life. Such arguments have been used to explain why humans approach a mortality plateau at very old ages (Missov & Vaupel 2015). If this type of selection bias occurs between the ages of 70 and 90, it could explain the chronological age-specific biases we observe in our study. Obviously, age perception adjusts over time, but it is less clear why raters would not be able to adjust to a selective bias as explained by frailty models. Such adjustment to the selective bias should happen for all cohorts and not just for specific cohorts.

Even though we find that the average perceived age approaches the average chronological age across all images rated, the power of perceived age as a biomarker of aging might rather be seen in its potential to explain deviances from chronological age. A biomarker of aging that only reveals chronological age is relatively uninformative, except for special cases when one does not know the chronological age of an individual as e.g. for a blood spot in a crime scene or for wild animals (Horvath 2013; Jarman et al. 2015). Perceived age holds great potential as a biomarker for biological age, because biomarkers of aging should inform about biological age differences beyond chronological age, and since perceived age is not well correlated to chronological age it shows the potential to provide exactly such additional information (Baker & Sprott 1988). This potential of perceived age as biomarker is confirmed by previous findings that show that perceived age corresponds to health measures, cognitive function and various hazard risks, and might be a better predictor of mortality hazard compared to chronological age (Christensen, Thinggaard, et al. 2009; Dykiert et al. 2012; Hwang et al. 2011; Belsky et al. 2015). Our main finding lays in the substantial shift that age perception underwent between 2002 and 2012, which was observed in almost all individuals (Fig. S3). This degree of generality is remarkable and suggests that the perception of biological age does not rely on specific characteristics only exhibited by a number of individuals, but rather that perceived age is a general and integrated biomarker of aging with the contemporary background population as the reference.

One of the riddles behind increased life expectancy is whether it occurs because aging is delayed — the deterioration in function starts at a later age in more recent cohorts — is slowed down, or individuals begin life at lower biological ages. A slow downed aging process, or reduced rate of aging, would be expressed in a slower deterioration process in more recent cohorts, which relates to the concept that the rate of aging changes over time (Vaupel 2010). If the biological aging process would be delayed and not slowed, we expect that such a delay would also apply to biological aging and should be reflected in perceived age. Hence, we would mainly expect to see a shift in the intercept in the deviance between chronological and perceived age between 2002 and 2012 (Fig. 2). We see a substantial shift in the mean perceived age that could suggest a shift as predicted by delayed aging or different starting levels of aging, but we also see a change in the slope of the deviance between 2002 and 2012 with respect to chronological age (Fig. 2 & Fig. S2). This reduction in slope is easiest associated with an expected change in the rate of aging. The challenge is that expectations regarding changes in rate of aging, delays of aging or different levels of biological age at birth are not mutually exclusive. Differences in starting levels of biological age have been found in cognitive function. Over the last century scores on standardized intelligence quotient (IQ) tests have increased — the so called Flynn effect (Flynn 1987). These improvements have been associated with better starting levels of cognition and support the preserved differentiation hypothesis (Salthouse 2006). However, efforts to track changes in cognitive performance have been largely focused on adults. Applying this concept to younger children might be difficult, and it might be difficult to distinguish between starting level differences and delayed degradation of cognition. Support for a delay in aging as a driver of increased lifespan has been revealed by investigating mortality rates but again no conclusive arguments have been made yet. Therefore the controversy continues about the process of aging and how prolonged lifespans in more recent cohorts have been achieved (Vaupel 2010).

We show that perceived age is likely a good and general biomarker of aging that can be used to track changes in biological rather than chronological age. Perceived age reflects past aging processes better than it predicts future outcomes of aging. This conclusion contrasts slightly with other studies that have shown how perceived age can improve predictions on the life prosperity of an individual; and these improvements were to a similar degree as predictions on life prosperity explained by chronological age (Christensen, Thinggaard, et al. 2009). Extending perceived age investigations to younger ages (<70), as suggested in the context of other biomarkers of aging (Belsky et al. 2015), would provide better means to shed light on a variety of controversial topics in demography and aging research, including whether increased life expectancy is achieved by delayed aging, by lower starting levels of biological age, or slowed aging. The advantage of a direct comparison of age perception shifts and changing life expectancy makes perceived age a very interesting and useful biomarker that is in addition quite cost efficient. However, this direct comparison also challenges us for interpretation of results. For instance, we would not be able to differentiate whether shifts in perceived age relate to changes in remaining cohort life expectancy or to changes in life expectancy at birth. We might want to aim for the use of biomarkers that go beyond qualitative or threshold levels to differentiate among biological age of individuals. Qualitative biomarkers that can be directly related to other well recorded shifts in demographic parameters might provide great potential for advancing our understanding of the aging process and its consequences across time — perceived age might be one of them.

## Methods

We captured facial images of 238 Danish women, and 144 Danish men aged 70+ (Fig. S1) in a standardized way between February 2001 and April 2001 (details in Christensen et al. 2009a). In November 2002, each of 20 raters assessed the perceived age of each individual in the 382 images without any knowledge of the participants chronological ages. In December 2012 this assessment was repeated on the same images by 9 raters (one rater was excluded as an outlier as this rater consistently rated much higher compared to the others). Here we only analyse average perceived age for each image over the 20 raters in 2002 or the 9 raters in 2012; hence we do not evaluate variance in perceived age among raters.

For the core part of the study, to address how perceived age has changed over time, we used linear models with difference between chronological and perceived age as response variable and chronological age, sex, and assessment year as fixed explanatory factors. We setup and ran models in the program R (R Core Team 2016) and selected among models based on Akaike’s Information Criterion (AIC), with a difference of >2 in AIC being considered to provide better support to the model with the lower AIC (Burnham & Anderson 2004). We evaluated model assumptions using diagnostic plots which suggested that a Gaussian error structure was appropriate for all models. To explore characteristics of the data and to detect potential biases in the data, we formulated additional models for which we used i) chronological age as response variable and sex as explanatory variable, ii) perceived age as response variable and sex and year as explanatory variables (Table S1), as well as iii) the difference in the deviance between chronological age and perceived age in 2002 and the deviance between chronological age and perceived age in 2012 as response variable, and sex and chronological age as explanatory fixed factors (Table S2).

A strength of our approach is that the variance in change in biological age over this ten-year period (2002-2012) is lower compared to the actual background population in 2002 and 2012, because we use the same images to evaluate the shift in age perception over the ten years. Such lowered variance increases the power of our analysis. In real life, some individuals might have for instance just gone through some severe illness when their image was taken. These individuals might appear relatively old but after some time of recovery they would have appeared to look younger again. Such variance is excluded in our approach because the images do not change, e.g. the signs of the illness are preserved, only the perception changed over the decade of investigation.

## Acknowledgements

We thank James Vaupel, Patrick Barks, and all members of the Max-Planck Odense Center on Biodemography for comments and discussions.

## Author Contributions

KC initiated and designed the study; UKS and LAL analysed the data; UKS wrote the first and final draft of the manuscript with substantial help from KC and LAL.

